# Incertohypothalamic A13 dopaminergic neurons are involved in fine forelimb movements but not reward

**DOI:** 10.1101/2022.03.30.486380

**Authors:** C Garau, J Hayes, G Chiacchierini, JE McCutcheon, J Apergis-Shoute

## Abstract

Tyrosine hydroxylase (TH)-containing neurons of the dopamine (DA) cell group A13 are well-positioned to impact known dopamine-related functions since their descending projections innervate target regions that regulate vigilance, sensorimotor integration and execution. Despite this known connectivity little is known regarding the functionality of A13-DA circuits. Using TH-specific loss-of-function methodology and techniques to monitor population activity in transgenic rats *in vivo* we investigated the contribution of A13-DA neurons in reward and movement-related actions. Our work demonstrates a role for A13-DA neurons in grasping and handling of objects that is independent from reward. A13-DA neurons respond strongly when animals grab and manipulate food items while their inactivation or degeneration prevents animals from successfully doing so - a deficit partially attributed to a reduction in grip strength. In contrast, there was no relation between A13-DA activity and food-seeking behavior when animals were tested on a reward-based task that did not include a reaching/grasping response. Moreover, motivation for food was unaffected as goal-directed behavior for food items was in general intact following A13 neuronal inactivation/degeneration. These results demonstrate a functional role for A13-DA neurons in prehensile actions that are uncoupled from reward and as such position A13-DA neurons into the functional framework regarding centrally-located DA populations and their ability to coordinate movement.

## Introduction

The degeneration of centrally-located dopamine (DA) neurons via disease or through experimental manipulation severely impacts the transformation of intent into action (Hefter et al., 1987; Oyanagi et al., 1989; Fearnley and Lees, 1991; Gerlach and Riederer, 1996; Chaudhuri et al., 2006; Jankovic, 2008). Classified centrally according to the cell group nomenclature A8 to A16 (Hokfelt, 1984; Bjorklund and Dunnett, 2007), the activity of spatially-distinct DA cell populations have been linked to the reinforcing properties of salient stimuli (Schultz et al., 1993; Fiorillo et al., 2003; Lak et al., 2014) and motor control (Panigrahi et al., 2015; Howard et al., 2017; da Silva et al., 2018; Hughes et al., 2020) that when coupled results in effort-based planning and execution that is critical for goal-directed behavior (Berridge, 2007; Grace et al., 2007; Kobayashi and Schultz, 2008; Palmiter, 2008). Although this framework is mostly supported by research focused on A9 ventral tegmental area (VTA) and A10 substantia nigra pars compacta (SNc) DA neurons, separate studies have demonstrated that other DA populations are also involved in motor and reward-based processes. For instance, activation of diencephalic A11-DA neurons in the posterior hypothalamus has been shown to modulate locomotor behavior potentially via their projections to the spinal cord (Koblinger et al., 2018). In contrast, A12-DA neurons located ventrally in the arcuate nucleus of the hypothalamus signal reward as it relates to food - a functional consequence likely mediated through their excitatory and inhibitory influence onto hunger and satiety-signalling cell populations, respectively (Zhang and van den Pol, 2016).

Anatomical work investigating their projection patterns suggests that A13-DA neurons may also be important for motor planning and/or motivation. Located in the rostromedial division of the zona incerta, A13-DA neurons contain dopamine (Negishi et al., 2020) and mostly send descending projections that innervate mesencephalic locomotor (MLR) regions linked to vigilance and locomotion (Sharma et al., 2018) as well as to the superior colliculus (SC) (Bolton et al., 2015) where sensory-motor integration related to fear-related defensive (Cohen and Castro-Alamancos, 2010; Zingg et al., 2017; Evans et al., 2018) and appetitive hunting/foraging behavior (Shang et al., 2019; Huang et al., 2021) occurs. Functional work on A13 subpopulations is, however, mostly lacking and as such their contribution to such processes is unresolved. Our study aims to investigate the functional properties of A13-DA neurons focusing on their potential role in motivation and motor regulation. To do so we have used transgenic TH-Cre rats in order to monitor and manipulate A13-DA neuronal activity while animals are engaged in reward and motor-based behavioural tasks. Our results indicate that A13-DA neurons are critical for skilled forelimb movements, specifically those involving prehensile action, that are independent of reward thereby revealing a novel DA system important for motor coordination and execution.

## Results

### A13-DA neuronal activity relates to forelimb movements that are independent of reward

To determine whether a relationship between A13-DA neuronal activity and motor and/or reward-based behavior exists we monitored A13-DA activity using fibre photometry in TH-Cre rats with the Ca^2+^ indicator GCaMP6s expressed in A13-DA neurons (Figure 1A). Rats were trained to lever press for access to a sipper containing 10% sucrose solution on a FR1 schedule of reinforcement. For temporally separating the motor component from the reward, a 5 sec delay was imposed between the lever press and sipper access. A chamber light indicated the onset of an active trial where the lever was extended and a response would be reinforced (Figure 1B1-top). We analyzed fibre photometry signals during three epochs - baseline, in response to lever press and during sipper availability - and a one-way repeated measures (RM) ANOVA revealed no statistical difference between the signals (F(2,14) = 1.738, P=0.21; n = 8; AUC: Baseline, −7.31 ± 5.23; Lever Press, 7.44 ± 5.31; Sipper, −10.0 ± 8.24) (Figure 1B1). When the operant requirement for sipper access was increased from one to three lever presses (FR1 to FR3) the motor responses coincided with a significant increase in A13-DA activity compared to both the baseline period and reward delivery (one-way RM ANOVA, F(2,10) = 6.384, P = 0.016; Tukey’s post-hoc differences: Lever Press vs. Baseline (P = 0.04) and Sipper (P = 0.02); n = 6; AUC: Baseline, −6.30 ± 3.44; Lever Press, 16.60 ± 11.53; Sipper, −9.06 ± 12.33) (Figure 1B2).

**Figure 1:**
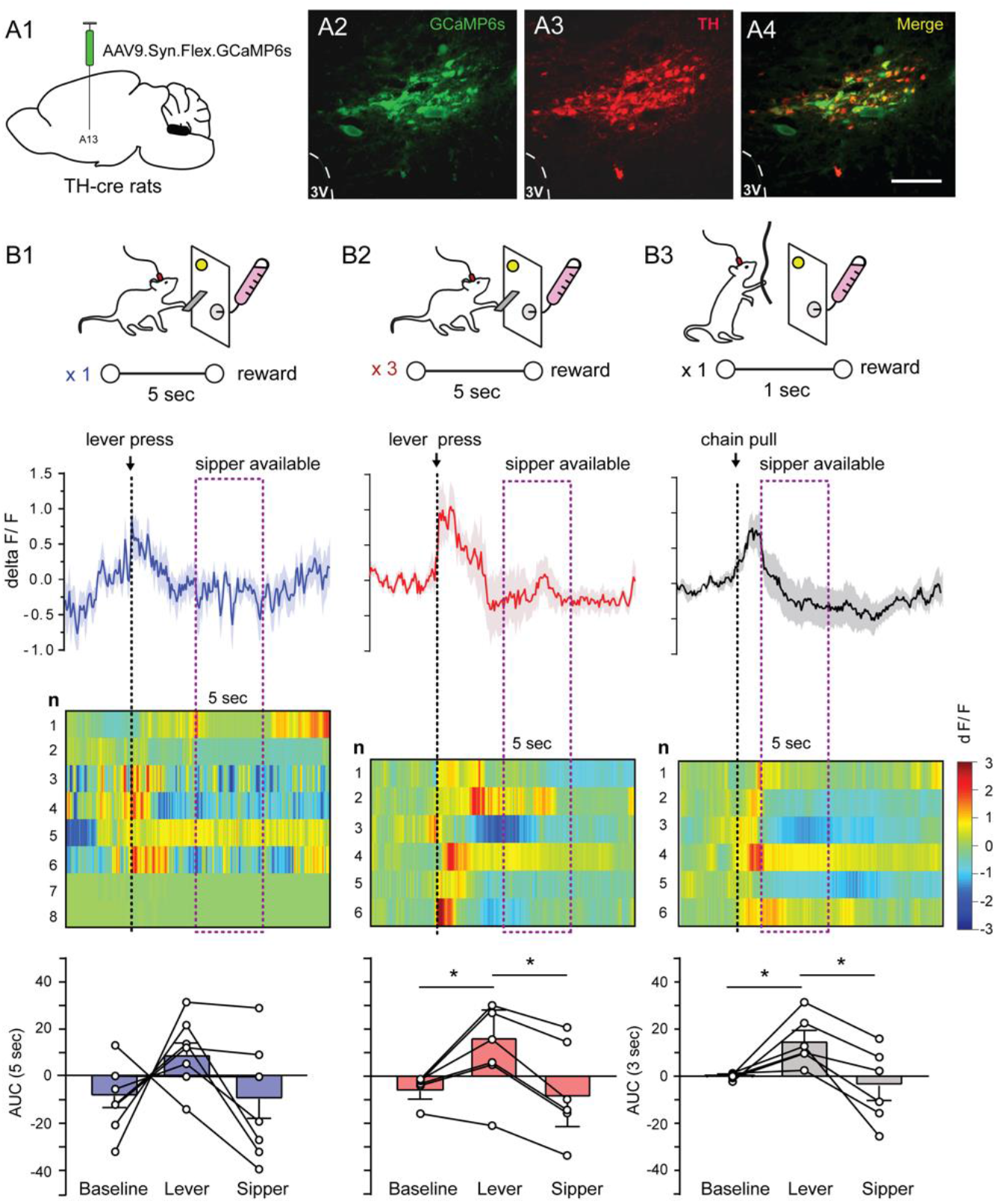
A13-DA population activity relates to operant forelimb movements. A1-A4, GCaMP6s expression in A13 TH-containing neurons of TH-Cre rats. A13-DA Ca^2+^ activity in response to lever press FR1 (B1) and FR3 (B2). Average photometry signal of lever press onset and sipper availability (top) followed by color plot depicting average of each rat (middle) and summary AUC data for baseline, lever press and sipper access (bottom). B3. Average photometry signal, response spectrogram for individual animals and summary data in related to chain pulling and sipper access. Scale bar, 50 μM; Abbreviations; 3V, 3^rd^ ventricle.

A13-DA responses were next related to a second operant task requiring forelimb movements where instead of lever pressing rats were trained to chain-pull for access to the same reward. Chain-pulling led to a similar significant increase in A13-DA activity compared to both baseline and sipper access (one-way RM ANOVA, F(1.43,7.15) = 7.336; P = 0.0235; Tukey’s post-hoc differences: Chain-pull vs. Baseline (P = 0.046) and Sipper (P = 0.01); n = 6; AUC: Baseline, −0.43 ± 0.56; Chain-pull, 15.21 ± 4.25; Sipper, 3.92 ± 6.40) (Figure 1B3). These findings demonstrate a link between A13-DA activity and an operant forelimb response.

Following learning, reward-based activity has been shown to shift from aligning with reward delivery to the stimulus predicting the reward (Schultz et al., 1993). It is thus unclear whether A13-DA photometry signals in the previous experiments were in response to the predictive light or the lever pressing. In an attempt to dissociate the two we monitored A13-DA activity while animals were active in a two-bottle choice task where rats were given intermittent access to lick from two bottles, each containing a solution with different rewarding properties (Figure 2A). A chamber light was again used to indicate an active trial which coincided with the extension of both sippers. Since rats were both signalled and given access to rewarding solutions without the need to lever press this task was therefore used to gauge reward-based conditioned and unconditioned responses that were independent of forelimb movements.

**Figure 2:**
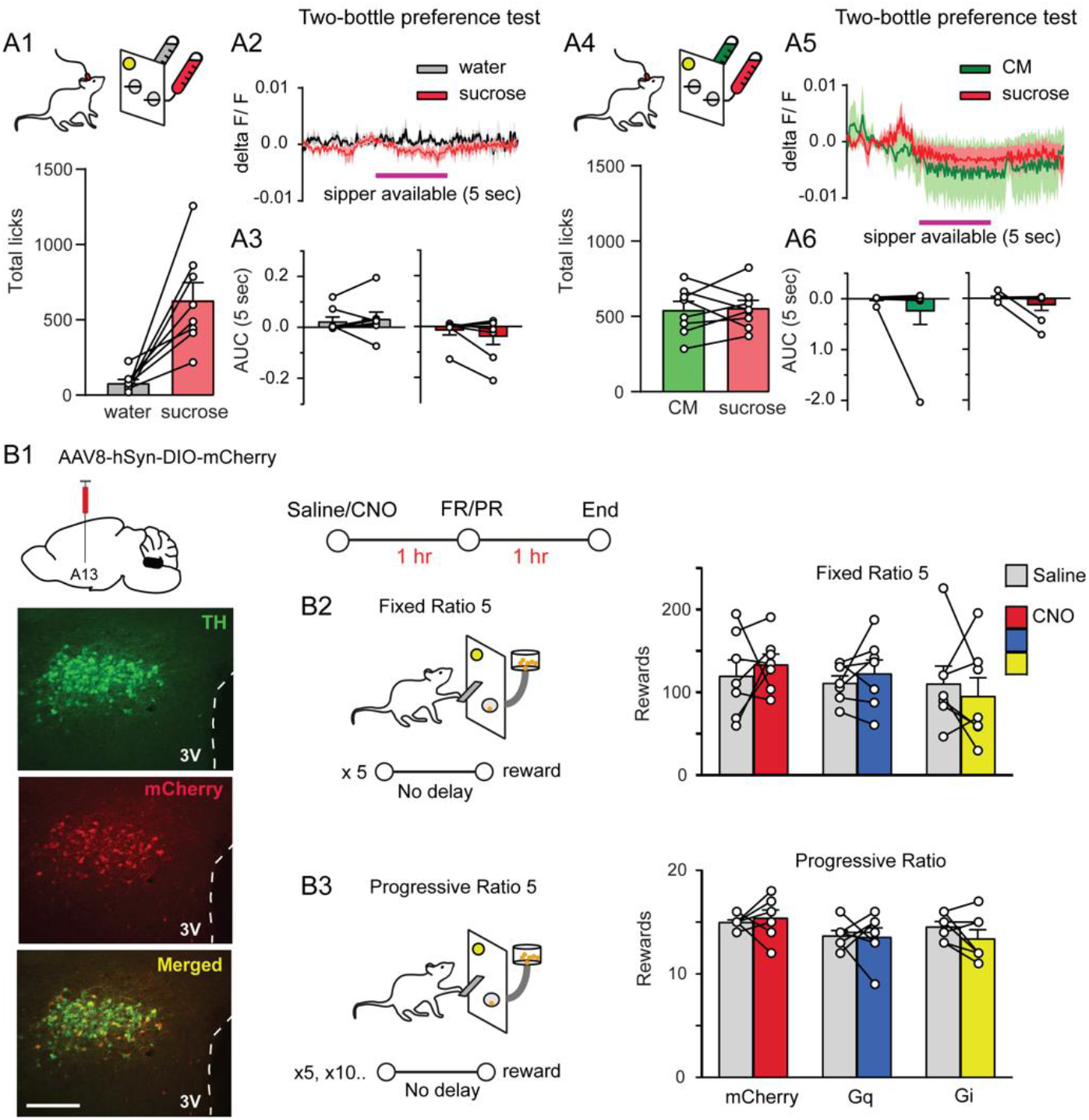
A13-DA activity is unrelated to motivated behavior towards reward delivery and consumption. In a two-bottle preference test A13 activity is unrelated to increases in licking for either a sucrose (A1-3) or condensed milk (CM) (A4-6) solution. A13-DA expression (B1) and manipulation of A13 activity did not affect motivation for a food reward on operant paradigms under FR5 (B2) or PR5 (B3) contingencies. Scale bar (B1), 75 μM: Abbreviations; 3V, 3^rd^ ventricle; AUC, area under the curve; CNO, Clozapine N-oxide; FR, fixed ratio; PR, progressive ratio.

When first given a choice between water and a 10% sucrose solution, as expected rats licked the sucrose-containing sipper significantly more than the one containing water (Paired TTest: T = 4.385, P = 0.0032; Total Licks: Water, 82.13 ± 22.77; Sucrose, 631.1 ± 115.2, n = 8) (Figure 2A1). When the corresponding photometry signals were analyzed there were no significant changes in A13-DA activity compared to baseline when licking for sucrose (Paired TTest: T = 1.35, P = 0.22; AUC: Sucrose, Baseline −0.02 ± 0.02; Sipper −0.04 ± 0.03; n = 8) or water (Paired TTest: T = 0.5477, P = 0.60; AUC: Water, Baseline 0.02 ± 0.01; Sipper, 0.03 ± 0.02; n = 8) (Figure 2A2,3). Similarly, when water was replaced with a more palatable condensed milk (CM) solution (50% in water) there was no change in activity for either the sucrose solution (Paired TTest: T = 1.56, P = 0.16; AUC: Sucrose, Baseline 0.02 ± 0.02; Sipper −0.14 ± 0.10; n = 8) or the CM (Paired TTest: T = 1.01, P = 0.35; CM, Baseline −0.01 ± 0.02; Sipper −0.25 ± 0.26; n = 8) (Figure 2A5,6) despite an increase in licking for CM that was comparable to that for sucrose (Paired TTest; T = 0.25, P = 0.81: Total Licks: CM, 544.0 ± 55.37; Sucrose, 556.4 ± 48.96; n = 8) (Figure 2A4). These results thus support the idea that A13-DA activity is unrelated to rewarding stimuli.

### A13-DA activity is not causally-related to lever pressing or motivation for a reward

We next tested whether A13-DA activity directly influences forelimb movements and/or motivation towards a reward. For this TH-re rats expressing designer receptors (DREADDs) used for manipulating A13-DA activity were tested on lever pressing behavioral tasks commonly used to investigate motivation (Arnold and Roberts, 1997). Expressing (Figure 2B1) and subsequently activating excitatory or inhibitory DREADDs through i.p. injections of the DREADD ligand CNO had no impact on either a FR5 or progressive ratio lever pressing task when compared to saline injections or to rats where only the reporter protein mCherry was expressed (FR5, 2-way ANOVA, Group: F(2,36) = 0.95, P = 0.40; Saline/CNO: F(1,36) = 0.06, P = 0.81; Interaction: F(2,36) = 0.43, P = 0.66) (Figure 2B2) (PR5, 2-way ANOVA, Group: F (2,36) = 3.21, P = 0.05; Saline/CNO: F(1,36) = 0.29, P = 0.60; Interaction: F(2,36) = 0.75, P = 0.48) (Figure 2B3). These results suggest that A13-DA activity is not required for reward-based motivation but neither is it necessary for effective lever pressing.

### Fine forelimb movements are related to A13-DA activity and dependent on an intact A13-DA system

Results from experiment 1 indicate that A13-DA neurons are active during a forelimb motor response but not so during reward delivery. This is in contrast to our loss-of-function tests where manipulating A13-DA activity did not impact lever presses to receive rewards. Since lever pressing can largely be accomplished by forelimb actions without fine control we reasoned that these seemingly conflicting results may be due to the fact that A13-DA activity may instead be related to paw/ digit movements. In order to test this scenario we trained TH-Cre rats on a skilled fine forelimb reaching and grasping task. Using a transparent Plexiglass chamber with a narrow opening at one end, rats were trained to reach through the opening to access a sucrose pellet. Following training, animals learned to successfully reach, grab and consume over 80% of pellets during a session (50 trials/session). Compared to baseline, A13-DA activity significantly increased when rats successfully reached for and grasped sucrose pellets (Paired TTest: T = 3.60, P = 0.009; AUC: Baseline, −0.09 ± 0.09; Pellet, 23.49 ± 6.58; n = 8) (Figure 3A1). When unsuccessful attempts were next analyzed, no increase in A13-DA activity was seen (Paired TTest: T = 0.26, P = 0.8; AUC: Baseline, −0.02 ± 0.06; Pellet, 0.90 ± 3.44; n = 8) (Figure 3A2) suggesting a role for A13-DA neurons in grasping that is independent of forelimb reaching movements.

**Figure 3:**
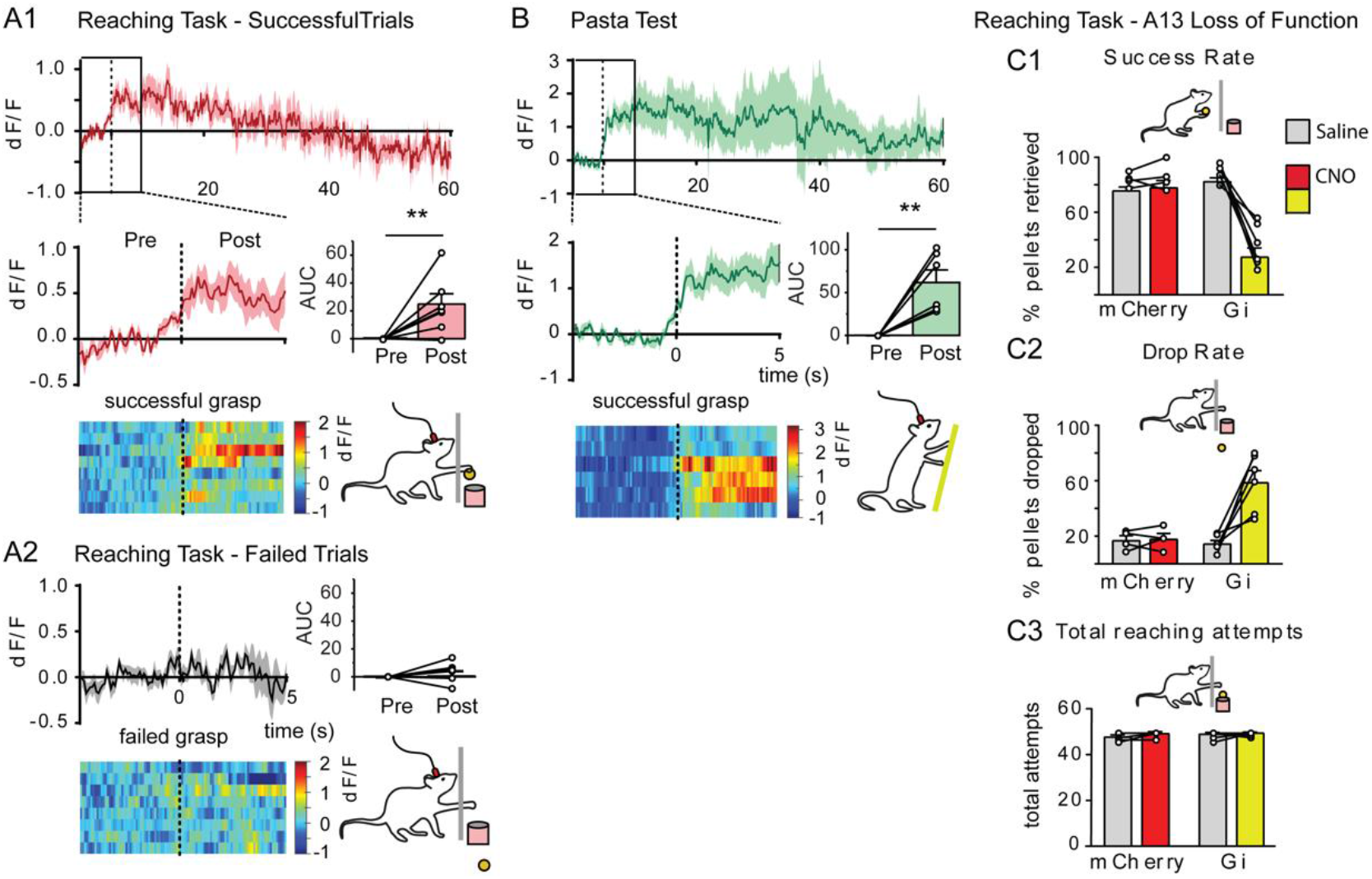
A13-DA activity is related to and is necessary for fine forelimb actions. A. Compared to baseline, A13-DA population activity increased with successful grasping and handling of sucrose pellets (A1) while failed attempts were unrelated to A13-DA activity (A2). B. A13-DA activity was elevated while rats handled and manipulated a dried spaghetti fragment. Pellet retrieval success rate decreased with A13-DA neuronal inactivation (C1) while the total amount of pellets dropped increased (C2). C3, No change in total reaching attempts was seen following A13-DA inactivation. Abbreviations: AUC, area under the curve; CNO, Clozapine N-oxide.

To further shed light on the relationship between A13-DA activity and fine forelimb movements A13-DA photometry signals were measured while rats handled and manipulated dried vermicelli fragments. This task is known as the pasta handling test and it requires no learning since rats naturally use their paws and fingers to guide a dried vermicelli fragment into their mouth for consumption. Consistent with the reaching task results, A13-DA activity increased when rats picked up and handled a dried vermicelli fragment (Paired TTest: T = 4.45, P = 0.007; AUC: Baseline, −0.37 ± 0.07; Handling, 62.30 ± 14.11; n = 6) (Figure 3B). The activity was similarly elevated throughout handling further supporting the idea that A13-DA neurons are important for fine forelimb movements.

Using a loss-of-function DREADD approach, we next tested whether A13-DA inhibition would impact the rats’ ability to successfully reach and grab sucrose pellets. Indeed, inactivating A13-DA neurons in Gi-expressing rats resulted in a significant decrease in the amount of successfully retrieved pellets when animals were injected with CNO compared to both CNO-injected mCherry and saline-injected Gi rats (2-way ANOVA, Group: F(1,8) = 26.20, P = 0.0009; Saline/CNO: F(1,8) = 24.27, P = 0.001; Interaction: F (1,8) = 28.56, P = 0.0007) (Figure 3C1). This retrieval deficit can be partially attributed to an inability to hold on the pellet once reached since CNO-injected Gi rats showed an increase in pellets dropped compared to control animals (2-way ANOVA, Group: F(1,8) = 12.11, P = 0.008; Saline/CNO: F(1,8) = 13.04, P = 0.007; Interaction: F(1,8) = 11.87, P = 0.009) (Figure 3C2). Interestingly, despite showing a strong deficit in retrieving sucrose pellets, CNO-injected Gi rats made a similar amount of reach attempts compared to control conditions demonstrating that motivation for the reward was intact (2-way ANOVA, Group: F(1,8) = 1.371, P = 0.28; Saline/CNO: F(1,8) = 2.23, P = 0.17; Interaction: F(1,8) = 0.56, P = 0.48) (Figure 3C3). To test whether these deficits were possibly due to changes in nociception or general motor deficits we next measured locomotor activity and exploration of Gi rats in an open-field chamber. Compared to controls, CNO-injected Gi rats showed neither differences in total distance traveled (2-way ANOVA, Group: F(1,8) = 0.34,P=0.58; Saline/CNO: F(1,8) = 0.25, P = 0.63; Interaction: F(1,8) = 0.12, P = 0.74) (Figure 4A) or time spent in the arena’s center (2-way ANOVA, Group: F(1,8) = 0.16, P = 0.70; Saline/CNO: F(1,8) = 0.015, P = 0.91; Interaction: F (1, 8) = 1.39 × 10^−6^, P = 0.99) (Figure 4B).

**Figure 4:**
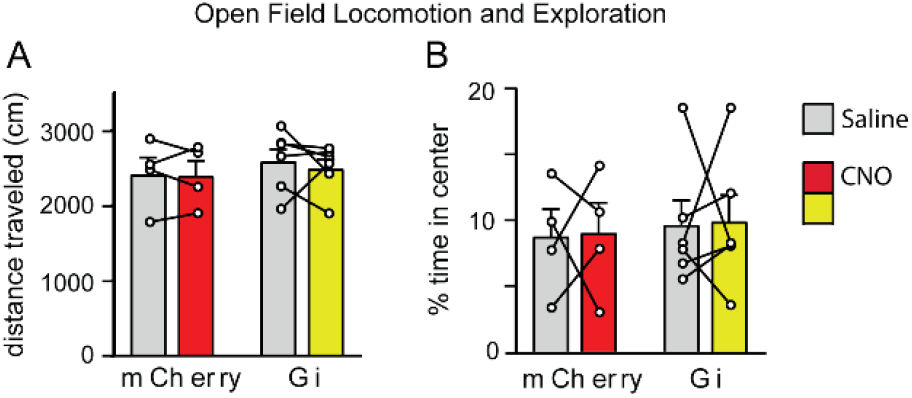
A13-DA inactivation does not impact exploratory behaviour. A. Compared to control, A13-DA inactivation did not affect the total distance traveled (A) or exploration of the middle area (B) of the open field. Abbreviation: CNO, Clozapine N-oxide.

### A13-DA neurons contribute to grip strength

The above results are based on behavioral tasks where a forelimb movement led to a reward. One interpretation of the results is that A13-DA activity may be encoding a successful movement-reward action. To investigate this possibility we tested rats on a grip strength test that was not associated with a food reward. Here, rats were required to hold onto a T-bar connected to a force meter while being gently pulled in the opposite direction. The amount of force rats could exert before letting go of the bar was then measured. CNO-injected Gi rats showed an approximately 60% decrease in grip strength compared to control conditions as measured in grams (2-way ANOVA, Group: F(1,8) = 1.63,P = 0.23; Saline/CNO: F(1,8) = 14.24, P = 0.004; Interaction: F(1,8) = 17.06, P = 0.003) (Figure 5A) but also when force was controlled for body weight (2-way ANOVA, Group: F(1,8) = 7.34, P = 0.03; Saline/CNO: F (1,8) = 17.30, P = 0.003; Interaction: F (1,8) = 16.91, P = 0.003) (Figure 5B). Taken together with the reaching and pasta-handling results, these data indicate that the contribution of A13-DA activity to fine forelimb motor movements is at least partially due to being involved in applying adequate levels of grip force for grasping actions.

**Figure 5:**
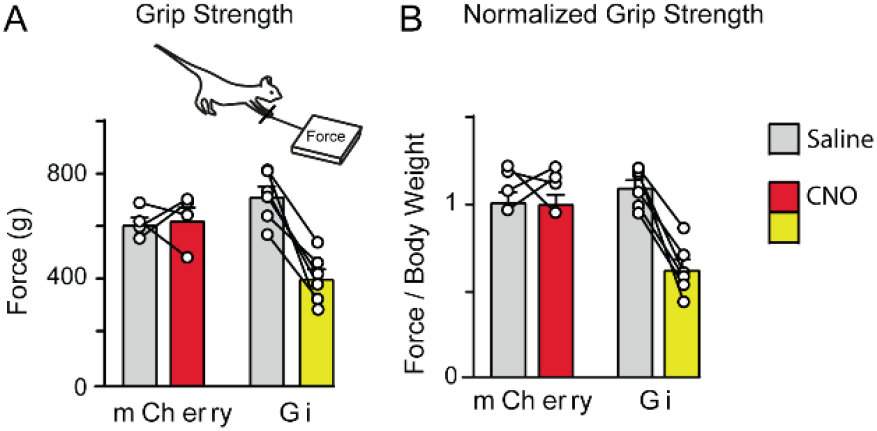
A13-DA inactivation led to a reduction in grip strength. Grip strength was significantly reduced following A13-DA inactivation when measured in total exerted Force (A) and when body weight was controlled for (B).

### A13-DA neurodegeneration severely disrupts pellet handling and grip strength

Dopamine cell degeneration linked to human diseases and their corresponding animal models has been shown to significantly disrupt grasping and handling behavior (Miklyaeva et al., 1994; Dunnett et al., 1998; Roberts et al., 2015; Thomas et al., 2017). To shed light on the potential contribution of A13-DA neurons to grasping and handling behavior in a neurodegenerative model, we next investigated the impact of A13-DA neuronal ablation on the reaching/grasping task. In a separate group of well-trained TH-Cre rats, a Cre-dependent viral construct expressing the apoptotic protein caspase-3 was injected into the A13. Pellet retrieval and drop-rate were monitored for 20 days post-surgery and pellet retrieval was seen to progressively decrease and plateau at approximately day 15 (Retrieval Rate: 2-way ANOVA, Group: F(1,11) = 57.51, P < 0.0001; Day: F(14,153) = 20.65, P < 0.0001; Interaction: F(14,153) = 15.42, P < 0.0001; Sidak post-hoc differences, Days 8-20) (Figure 6A1). Concurrently, pellet drop-rate increased across a similar time course (Drop Rate: 2-way ANOVA, Group: F(1,11) = 13.18, P = 0.004; Day: F(14, 153) = 5.26, P < 0.0001; Interaction: F(14, 153) = 4.02, P < 0.0001; Sidak post-hoc differences, Days 9-20) (Figure 6A2). Together, these findings demonstrate a striking deficit on a well-learned pellet handling and retrieval task following A13-DA neurodegeneration. Furthermore, measurements for total reach attempts revealed an interaction between group and days which post-hoc comparisons revealed was due to a significant difference on the first test day following surgery. (2-way ANOVA, Group: F(1,11) = 0.21, P = 0.65; Day: F(14,153) = 1.68, P = 0.07; Interaction: F(14,153) = 1.97, P = 0.024; Sidak post-hoc difference, Day 1) (Figure 6C). On all other days there was no significant differences. When next measuring grip strength, A13-DA ablated caspase rats showed comparable results to those measured in CNO-injected Gi rats where grip strength significantly decreased and plateaued a week following injection to approximately 60% of baseline (Force: 2-way ANOVA, Group: F(1,11) = 88.73, P < 0.0001; Day: F(15,164) = 1.910, P = 0.025; Interaction: F(15,164) = 6.014, P < 0.0001) (Force/BW: 2-way ANOVA, Group: F(1,6) = 22.40, P = 0.003; Day: F(15,90) = 2.104, P=0.016; Interaction: F (15,73) = 5.75, P < 0.0001) (Figure 6B).

**Figure 5:**
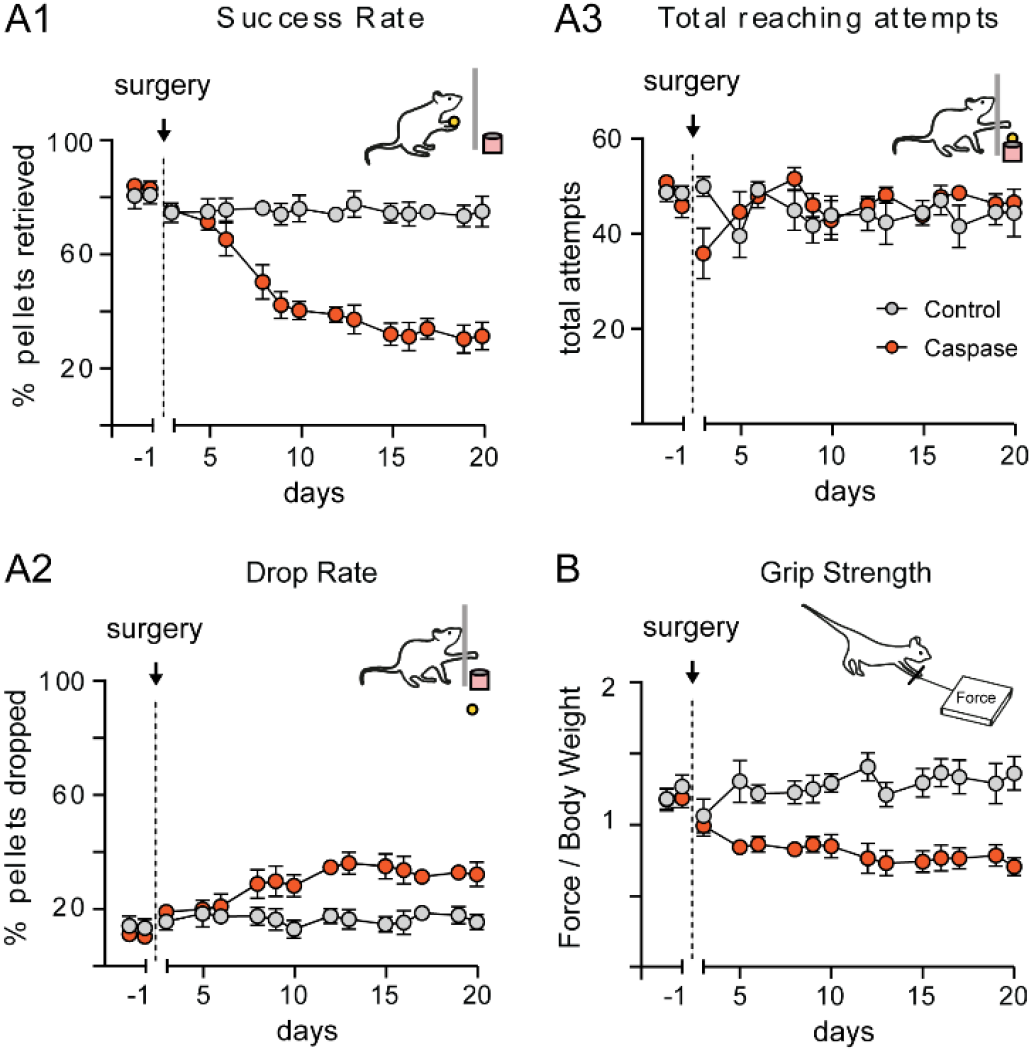
A13-DA neurodegeneration adversely impacted fine forelimb motor abilities. When tested on the reaching/grasping task, A13-DA caspase-mediated ablation led to a reduction in % of pellets retrieved (A1) and increase in pellets dropped (A2) while having no impact on total reaching attempts (A3). B. Grip strength as a function of body weight was significantly reduced following A13-DA ablation.

## Discussion

The above findings indicate that A13-DA activity relates to and is necessary for grasping and handling of objects. Motor forelimb actions such as lever pressing and chain pulling were tightly linked to A13-DA population activity as were skilled learned and unlearned grasping and handling movements. Consumption of palatable liquid rewards showed no relation to A13-DA activity and manipulations of A13-DA circuits did not impact motivation to lever press for rewards. In fact, despite the severe deficits in successfully grabbing and retrieving sucrose pellets caused by A13-DA neuro-inactivation or degeneration motivation for rewards was intact. The diminished ability to successfully retrieve pellets coincided instead with an increase in pellets dropped and a reduction in grip strength indicating that A13-DA activity is necessary for controlling paw and finger movements that are largely associated with prehensile actions.

### A13-DA neurons encode for prehensile actions that are uncoupled from reward

In rodents, skilled forelimb movements (SFMs) constitute a series of precisely timed sub-movements where the forelimb outwardly extends and ends with the paw and digits enclosing around a target object. DA activity is well-known to be critical for the planning and execution of complex motor sequences, including SFMs (WHISHAW et al., 1986; Bova et al., 2020), and has been shown to contribute in varying degrees to the different sub-components that make up a coordinated motor response. For instance, dopamine activity has been associated with action planning and initiation (Owen et al., 1995; Syed et al., 2016; da Silva et al., 2018), movement velocity (Panigrahi et al., 2015; Lofredi et al., 2018), force (Park et al., 2014; Hughes et al., 2020), locomotion (Kelly et al., 2012; Koblinger et al., 2018) as well as for the acquisition of learned skilled movements (Molina-Luna et al., 2009; Kawashima et al., 2012). Such a multi-faceted involvement for DA in movements consequently makes it difficult to target the exact contribution of DA activity on SFMs especially since separate DA populations have been shown to impact different SFM sub-movements through dissimilar pathways (Miklyaeva et al., 1994; Barneoud et al., 1995; Hosp et al., 2011; Bova et al., 2020). Our findings are the first to show a specific involvement of the A13-DA neuronal population in SFMs. In addition to disruptions in prehensile actions, increases in A13-DA activity were time-locked to grasping and were maintained while animals handled either a sucrose pellet or pasta fragment indicating that this dopaminergic population plays a specialized role in executing and maintaining grasping/handling movements. Our analyses however were unable to delineate which, if any in particular, prehensile sub-movements are encoded by A13-DA activity except that grip strength was severely impacted by A13-DA disruption. A future more systematic analysis of the underlying sub-movements and A13-DA activity will help reveal the precise A13-DA contribution to SFMs.

Prominent theories of dopamine function are based on the principle that dopamine-related motor functions and motivation are functionally coupled so that organisms can effectively plan, initiate, and execute motor programs for realizing a goal (Berridge, 2007; Niv et al., 2007; Salamone et al., 2018). This idea is highlighted by Mazzoni et al., (Mazzoni et al., 2007) where bradykinesia - a dopamine-dependent slowing of motor movements - in PD patients was shown to be at least partially attributed to a reluctance to move quickly rather than a deficit in motor speed or accuracy. This is in line with other work showing that the amount of vigor - defined as the speed, frequency, and amplitude of movements (Dudman and Krakauer, 2016) - at the onset of a planned action is strongly related to dopamine activity and determined by the predictability or incentive value of the goal (Pekny et al., 2015; Hamid et al., 2016; Bogacz, 2020; Hughes et al., 2020; Michely et al., 2020). Our results indicate that movement-related A13-DA activity does not encode for reward magnitude or reward consumption. Furthermore, it is unlikely to be associated with vigor since A13-DA activity was related to paw actions which occurred at the end of reach-to-grasp movements and unrelated to similar movements that ended in failed reaching attempts. Moreover, inactivation/neurodegeneration of A13-DA neurons had no impact on lever pressing or total reaching attempts indicating that motivation and, as a consequence, motor planning and initiation was intact. These results demonstrate a role for A13-DA neurons in SFMs that is uncoupled from the motivational factors that drive specific prehensile actions.

### Descending A13 projections innervate sensorimotor regions in the midbrain

Although their inputs are unknown, the functionality of A13-DA neurons can be inferred by considering our findings in light of their known efferent projections. A13 target areas include the periaqueductal grey (PAG) (Messanvi et al., 2013) and MLR (Sharma et al., 2018), both of which are regions composed of specialized networks that have been shown to shape sensory information into unique motor outputs (Cohen and Castro-Alamancos, 2010; Le Ray et al., 2011).The MLR for example is comprised of various subnuclei that together coordinate locomotor activity by influencing posture and gait (Skinner and Garcia-Rill, 1984; Josset et al., 2018). PAG activity similarly impacts muscle tension and coordinates motion but more so related to defensive behaviors such as freezing or fleeing in response to threats (Tovote et al., 2016). Our results, however, show that A13-DA neuron inactivation does not significantly impact exploration of a potentially threatening novel context or locomotor activity in general indicating that such motor actions are minimally affected by A13 inputs to these regions.

The A13 has also been shown to send axons to the SC (Bolton et al., 2015). The SC is a midbrain region important for coordinating movements towards or away from salient stimuli (Evans et al., 2018; Shang et al., 2019) and has been shown to promote orienting responses by, among other mechanisms, transforming visual and somatosensory information into saccadic responses that align with head rotations so as to stabilize gaze (Klier et al., 2001; Zahler et al., 2021). Interestingly, in both human and non-human primates, SC circuits have been shown to also encode for reaching movements (Himmelbach et al., 2013; Song and McPeek, 2015). In one study, neurons in the rostral pole of the primate SC were active during reaching movements and increased in activity when combined with visual input suggesting that specific SC circuits may be involved in gaze anchoring for the purpose of reaching and grasping target objects (Reyes-Puerta et al., 2010). Consistent with this, SC microcircuits in mice have been shown to be important in predatory hunting where a well-orchestrated repertoire of eye, head, body and forelimb movements are necessary for effective hunting - the end result being a well-timed reaching and grasping attack (Shang et al., 2019). Recent circuit mapping work has demonstrated that ZI circuits are also necessary for such prey-directed movements revealing a potential local A13-DA contribution to prey-grasping (Zhao et al., 2019) that in conjunction with SC sensorimotor responses may help coordinate effective predatory hunting (Shang et al., 2019).

### Functional architecture of brainstem-mediated skilled forelimb movements

SFM motor sequences have been shown to be generated cortically (Turella and Lingnau, 2014) and relayed to the spinal cord through direct descending cortico-spinal projections and indirectly via the reticular formation (RF), where pathways diverge innervating spinal circuits via the tectospinal, rubrospinal and reticulospinal tracts (Alstermark and Isa, 2012). Recent work has shown that functional divisions exist within the RF where separate subregions regulate distinct SFM motor components. For example, when tested on a reaching task, mice whose lateral RF glutamatergic cells were ablated showed deficits in reaching such that they consistently over-reached, missing the end-point (Ruder et al., 2021). Conversely, selective degeneration of medial RF glutamatergic neurons specifically affected pellet grasping, leaving reaching behavior intact (Esposito et al., 2014). Grip strength was unaffected in either study suggesting that these RF subregions are important for coordinating SFMs rather than generating the appropriate force necessary for grasping target objects. Our results show a clear relation between A13 activity and object grasping/handling which when disrupted resulted in reaching deficits that are at least partially due to an attenuation in grip strength. Interestingly, a recent retrograde tracing study has shown that A13 neurons innervate the RF, specifically the gigantocellular nucleus, thus revealing an anatomical substrate for A13 influence on RF output (Sharma et al., 2018). Whether or not the A13 also sends axons to adjacent SFM-regulating nuclei and what impact A13-RF projections have on SFMs are currently unresolved questions and important future research in determining central SFM networks.

### Dopaminergic activity is important for reaching and grasping movements

Reach-to-grasp actions are well-known to be symptomatic of DA neurodegenerative diseases and of their corresponding animal models (Miklyaeva et al., 1994; Olsson et al., 1995; Majsak et al., 1998; Whishaw et al., 2002; Pessiglione et al., 2003; Jo et al., 2015). For instance, detailed kinematic analyses of reach-to-grasp motions in Parkinson’s (PD) and Huntington’s (HD) patients have shown greater trial-to-trial variability in reaching for and grasping a target object compared to healthy individuals (Fellows et al., 1997; Alberts et al., 2000; Gordon et al., 2000; Lu et al., 2010). These deficits may result from an inability to shape appropriate anticipatory hand movements (Schettino et al., 2006; Rand et al., 2005; Lu et al., 2010) and also, in some cases, from alterations in the ability to apply the appropriate force (Stelmach and Worringham, 1988; Fellows et al., 1997; Reilmann et al., 2001; Guo et al., 2004) (but see also (Jordan et al., 1992; Nowak and Hermsdörfer, 2002)). Similar deficits in grasping have recently been linked to PD disease progression (Roberts et al., 2015) and, from a therapeutic standpoint, have been shown to be an effective diagnostic predictors of PD in early stages of the disease (Kandaswamy et al., 2018; Saikia et al., 2018). Surprisingly, pharmacotherapies for treating PD have shown to be only partially effective in reversing PD-SFM symptomology (Schettino et al., 2006; Tunik et al., 2007). For example, an analysis of hand kinematics in PD patients on/off L-Dopa therapy showed that reaching actions in medicated patients were only slightly improved while improvements in grasping were negligible (Negrotti et al., 2005). On the other hand, surgical therapies such as deep brain stimulation of the subthalamic nucleus – a region in close proximity to the A13 - show significant benefits in remediating both PD reaching and grasping deficits (Dafotakis et al., 2008; Pötter-Nerger et al., 2013). Such results suggest that separate mechanisms are at play for the different phases of SFMs that are circuit-specific and/or dopamine-dependent.

Hand actions are critically important in our daily lives. Everyday tasks such as cooking, eating, dressing and even holding playing cards are severely impacted in PD patients (Sjödahl Hammarlund et al., 2018) which consequently affects their physical and mental wellbeing (Kuopio et al., 2000; Dissanayaka et al., 2010). Our results demonstrate a clear and important role for A13-DA neurons in prehensile actions. From our analysis it is unclear, however, what signals are encoded by A13-DA circuits, the mechanisms by which dopamine or A13-DA co-transmitters might impact target regions and whether this distinct central DA population is affected in PD and HD. As such, future research investigating the contribution of this DA population to motor coordination and execution will be critical for positioning the A13 within the functional framework of central dopamine motor circuits and their relation to DA neurodegenerative diseases.

## Methods

### Animals and Husbandry

All male Long-Evans TH-Cre rats (450g-650g) that were used in this study were housed in pairs in individually ventilated cages under temperature-controlled conditions (21°C ± 2°C; 40%–50% humidity) and kept under 12 hr light/dark cycle, with lights on at 07:00. Post-surgery, rats were housed with bedding materials recommended by the NC3Rs and never single housed. Chow and water were available ad libitum except for a brief period of food restriction (12 hr before the first training session) and during experimental sessions. The animals were gradually food deprived so that their body weights were maintained at 85 ~ 90% of their ad libitum body weights throughout the experiments.

All procedures were carried out under the appropriate UK government Home Office license authority (PPL number: PFACC16E2) in accordance with the Animals [Scientific Procedures] Act (1986).

### Virus injection and implant surgery

Rats were deeply anaesthetised using isoflurane (5% / 2 L/min for induction, 2% for maintenance). The head was shaved and pre-operative analgesia was administered (bupivacaine 150 μl at incision site and meloxicam 1 mg/kg sub-cutaneous). Rats were mounted in a stereotaxic frame (David Kopf Instruments) and a thermostatic blanket was used to maintain a stable body temperature throughout surgery (37–38°C). Constant monitoring of oxygen saturation and heart rate was performed with a pulse oximeter. A scalp incision was made and a hole was drilled above the A13 for virus injection and fibre implantation (A13 from Bregma AP −2.5; ML ±1.3; DV −7.4). In addition, holes were drilled anterior and posterior to attach four anchor screws. A virus injection needle (10 μl Hamilton Syringe) was lowered into the A13 (DV −7.4 mm, from dura) and 1 μl of virus was injected over 10 min using a syringe pump (11 Elite, Harvard Apparatus, CA). A fibre optic cannula was implanted at DV −7.3 mm (0.1 mm above the virus injection site) (ThorLabs CFM14L10, 400 μm, 0.39 NA, 10 mm, sterilised using ethylene oxide). A layer of radio-opaque dental cement was used to seal screws and cannula in place (C&B Super-Bond, Prestige Dental) and a headcap was formed using dental acrylic (DuraLay, Reliance Dental). Care was taken to leave approximately 5 mm of the ferrule protruding for coupling to optical patch cable for later photometry recordings. Rats were housed in pairs immediately following surgery and at least 3-4 weeks was allowed post-surgery for virus expression before testing began.

Cre-dependent viruses were injected bilaterally into the A13 (AP-2.5; ML±1.3; DV-7.4). For fibre photometry experiments, a virus expressing GCaMP6s (AAV9.Syn.GCaMP6s.WPRE.SV40, ~1.9 × 1013 GC/ml; Addgene # 100843) was injected unilaterally in the A13 while for DREADD experiments viruses for reversibly activating (AVV8-Syn-hM3D-Gq-mCherry; Addgene # 44361), inhibiting (AVV8-hSyn-hM4D-Gi-mCherry; Addgene # 44362) or used as a control (AAV8-hSyn-mCherry; (Addgene # 114472) were injected bilalterally. Similarly for caspase A13-DA ablation, AAV1-flex-taCasp3-TEVp(Caspase); Addgene # 44580) or a control virus AAV8-hsyn-DIO-GFP was injected bilaterally.

### Fiber photometry

Fibre photometry equipment was similar to that reported elsewhere (Lerner et al., 2015) and consisted of two fibre coupled light sources powered by LED drivers. A blue 470 nm LED (Thorlabs, M470F3) and violet 405 nm LED (Thorlabs, M405F1) were sinusoidally modulated at 211 and 539 Hz, respectively, and passed through filters (470 and 405 nm). Both light paths were directed, via dichroic mirrors positioned inside filter cubes (FMC4_AE(405)_E(460–490)_F(500–550)_S, Doric Lenses), through a fibre optic patch cord (MFP_400/460/LWMJ-0.48_3.5m_FCM_MF2.5, Doric Lenses). The patch cord was then mated, using a ceramic sleeve, to the implanted fibre optic cannula. Emitted fluorescence was collected via the same fibre, through the patch cord and focused onto a photoreceiver (#2151, Newport). A signal processor (RZ5P; Tucker Davis Technologies) and Synapse software (Tucker Davis Technologies) were used to control LEDs, to acquire the lowpass filtered signal (3 Hz), and to perform on-line demodulation of the signal, separating isosbestic and calcium-modulated responses. Demodulated signals were acquired at 1,017 Hz. Behavioural events, such as licks and distractors, were routed to this system as TTLs and acquired simultaneously.

### Histology

Immunohistochemistry was used to verify fibre implant sites in the VTA and to check the extent of regional viral expression. At the end of experiment, all rats were terminally anesthetised with isofluorane and sodium pentobarbital (5 ml/kg) before being transcardially perfused with ice-cold phosphate-buffered saline (PBS) followed by 4% paraformaldehyde (PFA). Following perfusion, brains were removed and placed in 4% PFA for 24 hr before being transferred to a cryoprotectant 30% sucrose solution in PBS for at least 3 days. 40 μm coronal sections were made using a freezing microtome and stored in 5% sucrose solution in PBS with 0.02% sodium azide until staining.

Free-floating sections were first incubated in blocking solution (3% normal goat serum, 3% normal donkey serum, 3% Triton-X in PBS) for 1 hr before incubation with primary antibodies (mouse anti-tyrosine hydroxylase 1:1000, AB152; rabbit anti-mCherry 1:1000, ab167453; chicken anti-GFP 1:1000, ab13970; Abcam, Cambridge, UK) in blocking solution at room temperature on a shaker for 18 hr. Next, sections were incubated with secondary antibodies with either donkey anti-mouse AlexaFluor 488 (1:1000), donkey anti-Rabbit 594 (1:1000) or goat anti-Chicken AlexaFluor 488 (1:1000), (Abcam, Cambridge, UK) in PBS for 120 min at room temperature. Sections were mounted on slides using Vector Shield hard-set mountant (Vector Labs, UK). Between steps, sections were washed three times with PBS for 5 min and gently agitated on a laboratory shaker. Slices were imaged using an epifluorescent microscope (Leica, UK) to determine fibre placement and virus expression with the Paxinos and Watson (2007) rat brain atlas used as reference.

### Statistical analyses

All photometry and behavioural data were extracted from TDT files and analysed using custom Python scripts. These extracted data included signals for both photometry streams (calcium-modulated and isosbestic light levels) and timings of events (i.e. licks, lever presses, chain pulls, grasping, etc). The calcium-modulated signal was corrected for artefacts and bleaching using the isosbestic signal as a reference (Lerner et al., 2015). Once data were extracted, the corrected photometry signal was aligned to individual events signaled by TTL pulses or determined offline using video recordings. Data was then analyzed offline using Graphpad Prism. All data were expressed as means and standard error of means (SEM). Alpha was set at P < .05, all significance tests were two-tailed, and Sidak or Tukey’s HSD was used to correct for multiple comparisons.

### Behavioural Paradigms

Lever pressing, chain pulling and two-bottle choice experiments took place in operant behaviour chambers (Med Associates, VT, USA; 25 cm × 32 cm × 25.5 cm) housed inside large sound attenuating chambers. The lever, the sipper and the cue light were located on the same wall within the chamber. For tests reinforced with a liquid reward, a grid floor, comprised of stainless steel rods, was used in conjunction with contact lickometers, to record individual licks as rats consumed solutions from a spout recessed 5–10 mm from the chamber wall. For tests reinforced with sugar pellets, one pellet was delivered into a food hopper for each successful response. In these tasks a light signalled the beginning of an active trial. For tests reinforced with a liquid reward, sippers were extended after successful responses and were made available for 5 sec when lick contact was made. If there was no contact sippers were retracted after 20 sec. For DREADD experiments, saline or Clozapine N-Oxide (CNO) at 1 mg/kg was injected 1 hour before the start of experiments.

#### Lever pressing task

Using a fixed ratio (FR) or reinforcement rats were trained to press 1, 3 or 5 times for a reward (sipper access or sucrose pellet). In the Progressive Ratio, increasing number of operant responses are needed for each successive reward (5, 10, 15, etc). A 5 second delay was imposed between the operant response and reward availability.

#### Two-Bottle Free Choice Task

Prior to testing, rats were habituated to the presence of two drinking bottles for 3 days in their home cage. Following this acclimation, two different experiments were performed where in each rats were given simultaneous intermittent access to two sippers each containing a different solution: either sucrose or water or sucrose and condensed milk (CM). The bottles were presented a total of 20 times each.

#### Chain Pulling

The rat chain-pulling test was performed as described in detail elsewhere (Bayley et al., 1996). Instrumental procedures were performed using a metal chain 0.8 cm in diameter and 16 cm in length, which was attached to electronic switches mounted on the chamber ceiling. Following a successful pull, rats licked the available sipper while simultaneously gripping the chain and so behaviourally the motor response was largely inseparable from the sipper access. We thus analysed 3 sec instead of 5 sec photometry segments so as to minimise overlap between the motor and reward period of the task.

#### Reaching task

A rodent skilled reaching task was used to evaluate forelimb motor function. Rodents were trained to reach out through a slot to grab and retrieve a sucrose pellet for them to consume. The single pellet reaching box was rectangular, constructed of clear Plexiglas whose dimensions are 25×25 cm, and 50 cm high and designed to allow filming of the reaching behaviour (Whishaw and Pellis, 1990). A vertical slot 10 mm wide and 10 cm high on one wall opened to an external shelf that was mounted 3 cm above the floor. The pellet was placed on an indentation on either side of the opening to prevent rats from lapping the food with their tongue. Because the hand pronates medially to grasp, this placement required use of the contralateral hand for pellet retrieval.

Food restricted rats were first habituated for as long as 1 week by placing them in the boxes for 10 min each day. Commercial food pellets weighing 45 mg were initially available on the cage floor within a tongue distance on the shelf to associate the cage and shelf position with a food reward. Pellets were incrementally distanced on the shelf until the animals were forced to reach with a hand to retrieve the food.

#### Pasta test

The pasta test used was based on (Tennant et al., 2010). Uncooked Skinner brand vermicelli pasta (1.5 mm diameter) were cut to 7 cm lengths using a razor blade. To overcome neophobic responses and to establish skill in handling, animals were homecage exposed to the pasta pieces between 5-10 days of exposure, 4 pieces each time, prior to testing. Rats were also similarly habituated to pasta in the testing chamber (3-5 sessions, 25×25 cm, and 50 cm Plexi-glass chamber).

#### Grip Strength

This test is based on the tendency of a rat to instinctively grasp a bar or a grid when suspended by the body and permits assessment of forelimb strength. The test apparatus (Grip Strength Meter, Ugo Basile, Italy) consisted of a grasping bar attached to a force transducer in order to measure the maximum force applied by the rat during the pull. The unit of force used is grams-of-force. Each animal was handled via the body and brought near the bar, allowing the grasping of the grid with both forepaws and then gently pulled back until they released it. Measurements were discarded when rats only used one paw, used its hind paws, turned backwards during the pull, or released the bar without resistance. Animals were trained and tested on two consecutive days, using the same protocol. Five such measurements were obtained for each animal, and the resting period between each pull was 1 min (Manfré et al., 2017). The three best of each session were averaged and used for analysis. Results were expressed as total force (gm) and as normalized by body weight (force in grams/body weight in grams).

### Locomotion & Exploration

Rats were tested in a brightly illuminated open arena made of opaque white Plexiglass (50 × 50 cm floor area). Each animal was placed gently on the central zone and its behaviors were recorded by a camera for 15 min. The neurobehavioral parameters were computed offline at a rate of 30 frames/s using a video-tracking software (ANY-maze, Stoelting, CO, USA). Briefly, total distance traveled and time spent in the center and bordering regions was quantified by automated video-tracking and the percentage of time spent in the center of the arena was calculated. Tests were conducted between 1000 and 1500h. In order to maintain arena novelty, each rat was tested with a frequency of once a week and the context was adjusted by adding local and distal cues (i.e. striped walls).

## Acknowledgements

The authors acknowledge the help and support from the staff of the Division of Biomedical Services, Preclinical Research Facility, University of Leicester, for technical support and the care of experimental animals as well as colleagues in the department of neuroscience, psychology and behavior at the University of Leicester for their academic contribution. This work was funded by the Wellcome [grant #209023/Z/17/Z to J.A-S] and the Leverhulme Trust [grant #RPG-2017-417 to J.E.M. and J.A-S.].

